# hemispheR: an R package for fisheye canopy image analysis

**DOI:** 10.1101/2022.04.01.486683

**Authors:** Francesco Chianucci, Martin Macek

## Abstract

Hemispherical photography is a relevant tool to estimate canopy attributes such as leaf area index (LAI). Advancements in digital photography and image processing tools have supported long-lasting use of digital hemispherical photography (DHP). While some open-source tools have been made available for DHP, very few solutions have been made available in R programming packages, and none of these allows a full processing workflow to retrieve LAI and other canopy attributes from fisheye images.

To fill this gap, we developed an R package (*hemispheR*) to support the whole processing of DHP images in an automated, fast, and reproducible way. The package functions, which are designed for step-by-step single-image analysis, can be performed sequentially in a pipeline, while allowing inspecting the quality of each image processing step. The package allows to analyze both circular and fullframe fisheye images, collected either with upward facing (forest canopies) or downward facing (short canopies and crops) camera orientation. In addition, the package allows to implement two consolidated LAI methods (LAI-2000/2200 and 57° method).

A case study is presented to demonstrate the reliability of canopy attributes derived from *hemispheR* in temperate deciduous forests with variable canopy density and structure. Canopy attributes were validated against either results obtained from a reference proprietary software, either by benchmarking measurements obtained from terrestrial laser scanning. Results indicated *hemispheR* provide reliable openness and leaf area index values in forest canopies as compared with reference values.

By providing a simple, transparent, and flexible image processing procedure, *hemispheR* supported the use of DHP for routine measurements and monitoring of forest canopy attributes. Hosting the package in a Git repository will further support development of the package, through either collaborative coding or forking projects.

## 1. Introduction

The relevance of hemispherical photography for indirect canopy structure determination in forest monitoring has already been demonstrated (for a review, see Breda, 2003; Chianucci, 2019; Jonckheere et al., 2004; Yan et al., 2019). Hemispherical (also called fisheye) photography has been a pioneering method in forest ecology, with first applications dating back about seventy years ago (Anderson, 1964; Evans and Coombe, 1959). Since then, the method has been used for a wide range of applications, including phenology (Brown et al., 2020; Lang et al., 2017), ecosystem diversity (Sercu et al., 2017; Skelly et al., 2005), forest management (Chianucci et al., 2016; Thomas et al., 1999), microclimatology (Kašpar et al., 2021) and remote sensing (Fang et al., 2019). Among the features of hemispherical photography, a strong advantage of the method is that it can provide leaf area index (LAI) without knowledge of the foliage angle distribution (Chianucci and Cutini, 2012). In addition, the method provides the most comprehensive set of attributes related to canopy structure at the largest canopy footprint. The recent advancements in digital photography and image processing technology, combined with the low equipment costs compared with scientific optical instruments, have contributed to rising interest and widespread use of digital hemispherical photography (DHP) in forest ecology (Chianucci, 2019).

Notwithstanding these improvements, a great obstacle in the adoption of DHP is the sensitivity of results to image acquisition (Glatthorn and Beckschäfer, 2014; Macfarlane et al., 2000), and the complexity of fisheye image processing (Chianucci, 2019; Macfarlane, 2011). All these factors hinder the standardization and comparability between image acquisition and procedures adopted using hemispherical photography, while also limiting the accessibility of the method by non-experts (Chianucci, 2019). Therefore, an effective step towards standardizing image processing of DHP while making this method accessible to a wider audience is to develop an open-source tool to process these images reliably and efficiently.

To date, some free tools have been made available for hemispherical photography analysis, such as Gap Light Analyzer (GLA; Frazer et al., 1999), CAN-EYE (Weiss, Marie and Baret, 2017) and CIMES (Gonsamo et al., 2011), but none of these existing free solutions has been regarded as the ‘reference standard’ by the scientific and end-user community. In addition, only few tools have been made available in R to deal with canopy fisheye images. R is particularly interesting as it is both free and supported by a large and diverse user community, particularly through the development of packages – the “fundamental unit of shareable code” – which strongly supports open-science best practices (Atkins et al., 2022).

Currently, the existing R packages dealing with DHP focus on specific processing steps, such as thresholding (package ‘caiman’; Díaz et al., 2021)), gap fraction inversion (‘hemiphoto2LAI’; Zhao et al., 2019), canopy openness retrieval (‘Sky’; Bachelot, 2016). In addition, only ‘hemiPhot’ (ter Steege, 2018) allows inferring LAI from fisheye images, but this package has no flexibility in setting hemisphere zenith rings and azimuth sectors for the analysis of images. This calls for the development of a comprehensive and flexible R tool to perform all the steps required for processing digital hemispherical images in a single package.

In this contribution we introduced *hemispheR*, an R package to allow a step-by-step analysis of digital canopy images collected with fisheye lens. The next section illustrates the basic theory of the method, the digital image processing steps required to analyze fisheye images, and the package content and functioning. The reliability of canopy attributes derived from DHP images, processed using the *hemispheR* package, has been compared against canopy estimates derived from a reference proprietary software, and by benchmarking effective leaf area index values against reference values derived from terrestrial laser scanning.

## 2. Basic theory of hemispherical photography

DHP is based on digital images collected with the camera equipped with a wide-angle fisheye lens. In forest stands, images are acquired below-the canopy, with the camera oriented towards the zenith (upward-facing). In short canopies, such as agricultural crops, DHP is also used by orienting the camera downward (Liu and Pattey, 2010). Depending on the image field of view (FOV), principally, two types of images can be collected using the fisheye lens: (i) circular fisheye images contain the entire 180° hemisphere in a circle inscribed inside the rectangular image, while (ii) fullframe fisheye images reach a 180° FOV only across the diagonal (Figure 1). Compared with the circular format, fullframe images lack information about part of the canopy observed near the horizon from the sampling point, but on the other hand they have the advantage that all the image pixels are used to sample the canopy, which result in higher effective image resolution (number of pixels used to sample the canopy). However, analysis of fullframe images has been limited, as the available fisheye canopy tools have been mostly tailored for circular images only.

**Figure 1.**
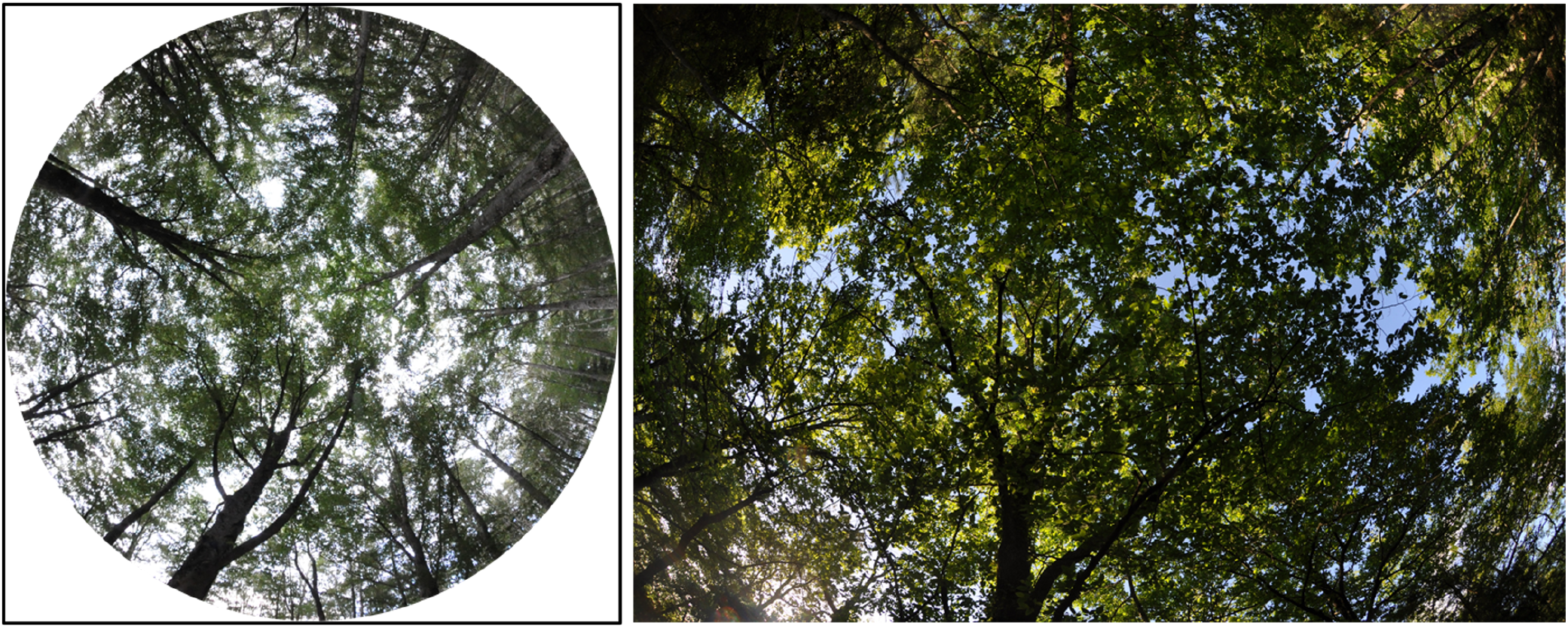
Circular (left) and fullframe (right) fisheye images of forest canopy.

The first step of DHP processing is classification into canopy and sky pixels, from which the gap fraction is calculated for a user-defined number of zenith rings and azimuth segments. LAI is derived from inversion of the Beer-Lambert law analogy for plant canopies:

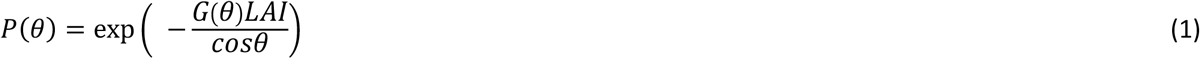

where *P(θ)* is the gap fraction, *G*(*θ)* is the *G*-function, which corresponds to the fraction of foliage projected on the plane normal to the *θ* direction, and *cosθ* accounts for the optical path length.

*The* G-function depends on leaf inclination angle and the leaf inclination angle distribution, which are generally *a-priori* unknown. In fisheye sensors, the influence of G-function on LAI is resolved by the hemispherical integration, as it gives (Miller, 1967):

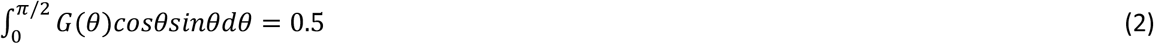

Hence, the inversion of angular gap fraction in fisheye methods, considering a discrete number of zenith rings, is:

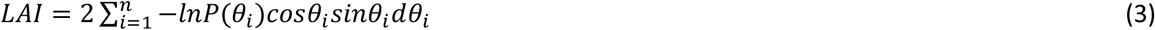

Where *n* is the number of *ith* zenith rings, *sinθ*_*i*_*dθ*_*i*_is the weigthing vector, which is normalized to sum as one. Alternatively, G-function can be calculated from gap fraction by assuming an Ellipsoidal leaf inclination angle distribution (Campbell, 1986), which is defined by a single parameter *x*, which is derived via an iterative method.

## 3. Overview of digital hemispherical image processing and hemispheR package

The key steps to analyze digital fisheye images are basically:

i. import an image channel and apply a circular mask (in case of circular fisheye images);
ii. apply a method to classify the single-channel image pixels into canopy and sky;
iii. divide the hemisphere into a number of concentric annuli (zenith rings) and radial sectors (azimuth segments) to retrieve the angular distribution of gap fraction;
iv. apply theoretical formulas relating gap fraction to canopy attributes.

Specific features need to be considered when dealing with fisheye images:

- circular mask parameters (x, y coordinates of the center and radius of the circle) are needed to exclude outer pixels from analysis;
- lens projection function needs to be considered to correct for fisheye lens distortion;

We also considered the following features relevant for fisheye image analysis:

- calculating vegetation indices (in case of downward looking images; Liu and Pattey, 2010);
- back-correcting the gamma function to unity (Macfarlane et al., 2007b, 2007a);
- enhance image contrast to ease thresholding (Macfarlane et al., 2014).

The *hemispheR* package uses the functionality of ‘raster’ package (Hijmans, 2021), which ensures faster processing of images not otherwise possible when dealing with other image formats in the R environment. The ‘raster’ package also has the strong advantage of allowing any kind of raster graphic image formats (i.e, pixel matrix), including raw imagery.

The package has been hosted in Gitlab at the following URL: https://gitlab.com/fchianucci/hemispheR. Gitlab, as well as Github, are increasingly used hosts for developing R packages, as it provides a simple and straightforward way to develop and install R packages (Decan et al., 2015).

The package features the following functions, which are ordered sequentially, following the fundamental image processing steps:

i. *import_fisheye()*: imports an image channel (or a mixing channel) and applies a circular mask (in case of circular images);
ii. *binarize_fisheye()*: thresholds the selected image channel and returns a binary image;
iii. *gapfrac_fisheye()*: calculate the gap fraction for defined zenith and azimuth bins;
iv. *canopy_fisheye()*: infer canopy attributes from the angular distribution of gap fraction Additional utility functions have been created to complement the package:
v. *camera_fisheye()*: returns the parameters needed for applying a circular mask for a known set of camera and lens models;
vi. *zonal_mask()*: divide the image into four sectors, which are used for zonal thresholding (useful in case of image collected in uneven light conditions);

All these functions are described in detail in the next paragraphs.

### 3.1 Import a fisheye image channel

The *import_fisheye()* function allows to import an hemispherical image in the R environment. The function requires selecting the image channel number using the ‘*channel’* argument, considering by default the blue one (which is set as 3 in RGB images). The blue channel is generally preferred in canopy image analysis, as it allows best contrast between canopy and sky, particularly when images are acquired in diffuse sky conditions. As an alternative to selecting a single image channel, the function can create a new single channel image by channel mixing with the following options available:

- Grenn Excess Index (Woebbecke et al., 1995): *GEI* = 2*G* − *R* − *B*
- Green Leaf Algorithm (Louhaichi et al., 2008): 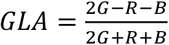
- *Luma* = 0.3*R* + 0.59*G* + 0.11*B* (standard for encoding analog to digital signals, close to visual perception of luminance of color image)
- 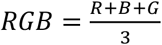 (average of RGB channels)
- 2*BG* = 2*B* − *G* (increases contrast between blue and green channels)

In case of GEI, GLA and 2BG, pixel values are then rescaled to 0-255 range for subsequent analyses. Both GEI and GLA have been included in the package to make it suitable also for downward looking imagery. However, some of these options can also be used in case of image collected in non-diffuse sky conditions, to deal with uneven illumination of the images (see an example in Figure *2*).

**Figure 2.**
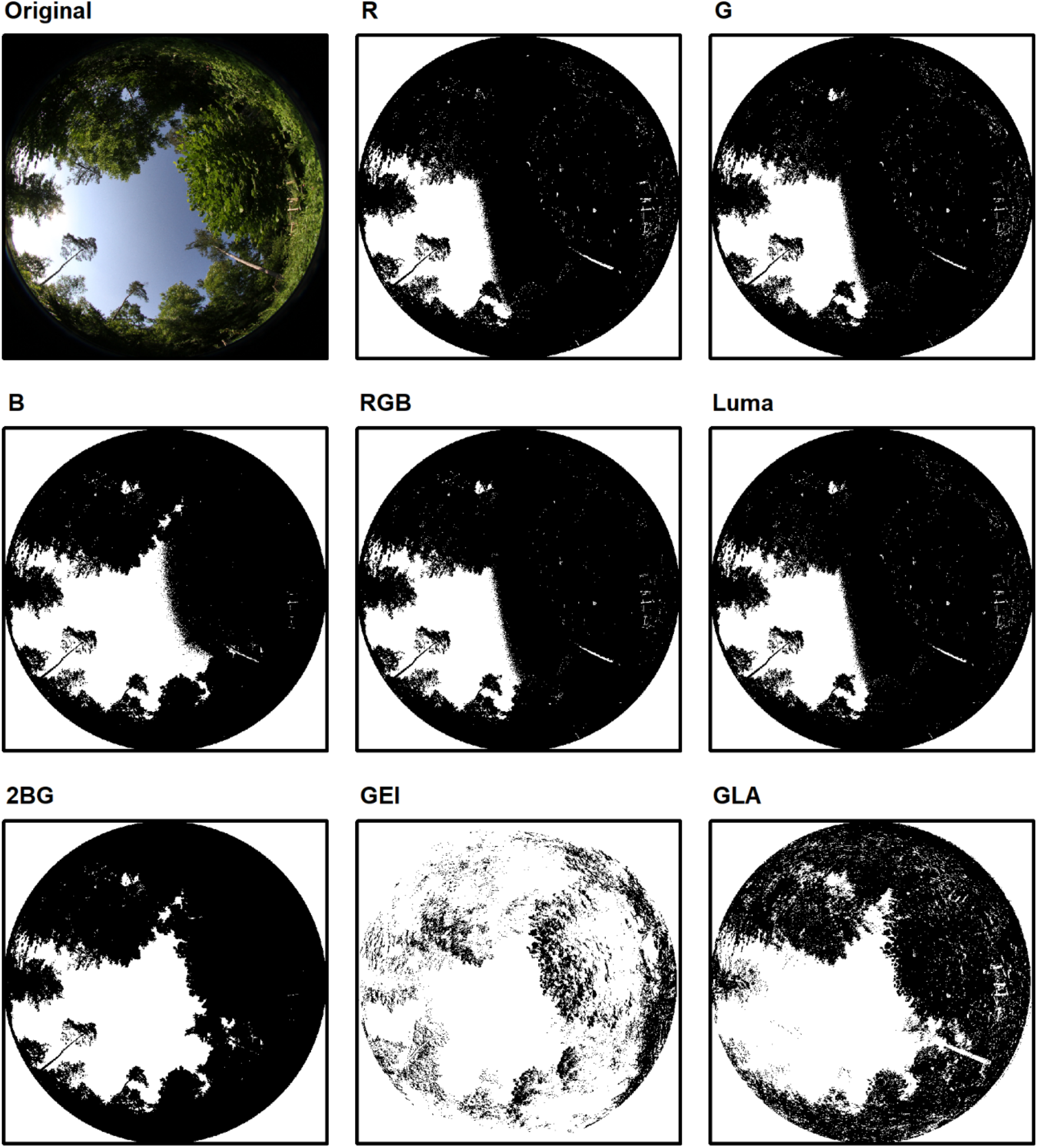
An example of a badly exposed image, with the scene lighted by direct sun from a low angle. In such situation, the channel selection is more influential for image thresholding; use of mixing channel options like 2BG or GLA can reduce the issues of wrong camera exposure. Images have been classified using ‘Otsu’ method in the *binarize_fisheye*() function.

Both circular and fullframe fisheye images can be imported from the function, using the argument ‘*circular*’. In case of circular images (circular=TRUE), the mask parameters (xc, yc, rc) need to be set to apply the circular mask, using the ‘*circ*.*mask’* argument. These parameters can be also derived from a list of available camera + fisheye lens sets, using the *camera_fisheye()* function (Table *1*). If omitted, the mask is created automatically in circular images.

**Table 1.**
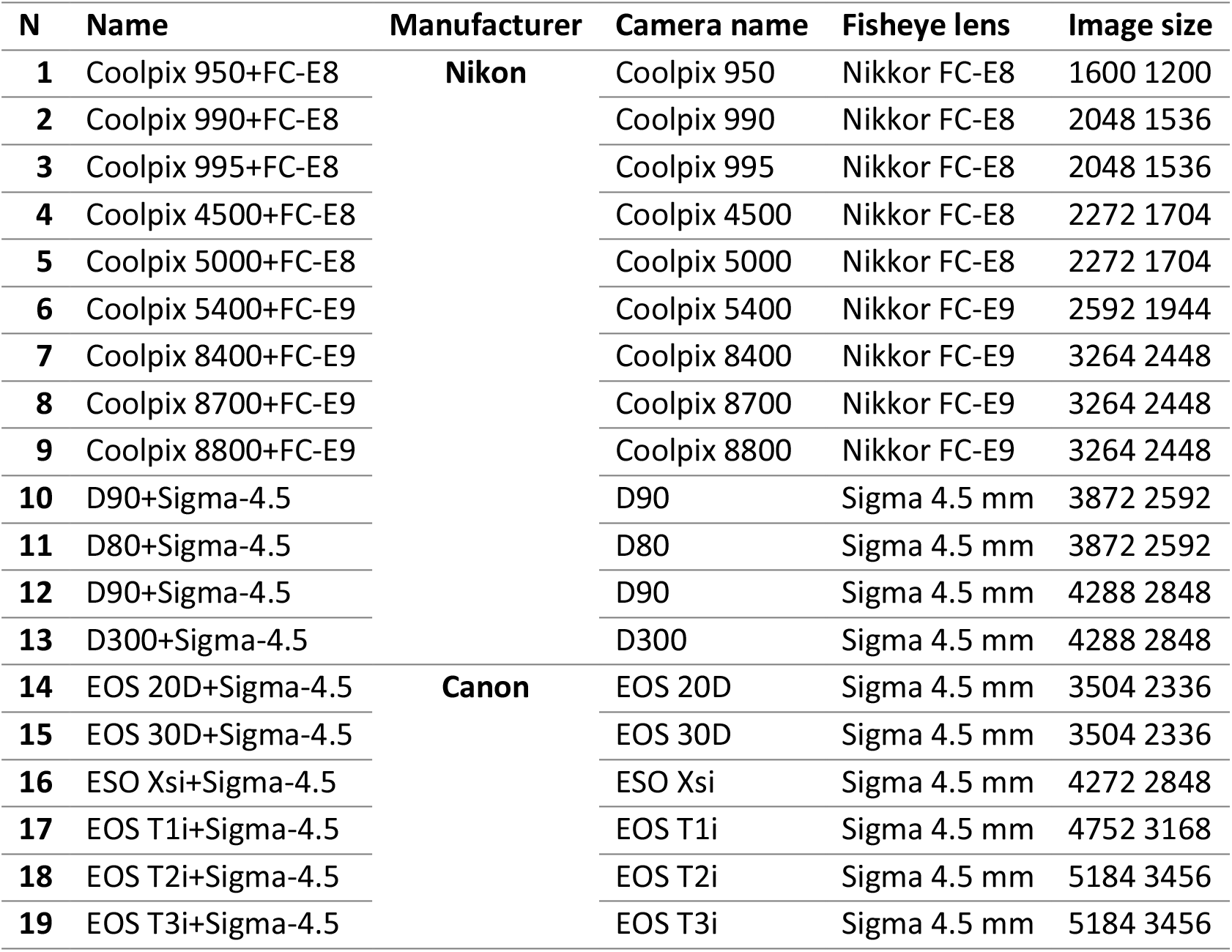
List of available camera and lens equipment for setting a circular mask using the camera_fisheye() function. The list can be screened in R by typing ‘*list*.*cameras’*.

The ‘*gamma’* argument allows to define the input image gamma, which is then back-corrected to unity (default value is set to 2.2, as typical of most image formats). Adjusting the gamma is strongly recommended in fisheye images, as it affects the calculation of gap fraction (Macfarlane et al., 2007). Finally, the ‘*stretch’* argument allows to apply a linear contrast stretch, by saturating 1% of pixels at the dark and light ends of the brightness histogram (Macfarlane et al., 2014). Finally, two optional arguments ‘*message*’ and ‘*display’* allow respectively to print out the mask applied to the image, and plot the imported image channel, along with the circular mask applied (Figure *3*).

**Figure 3.**
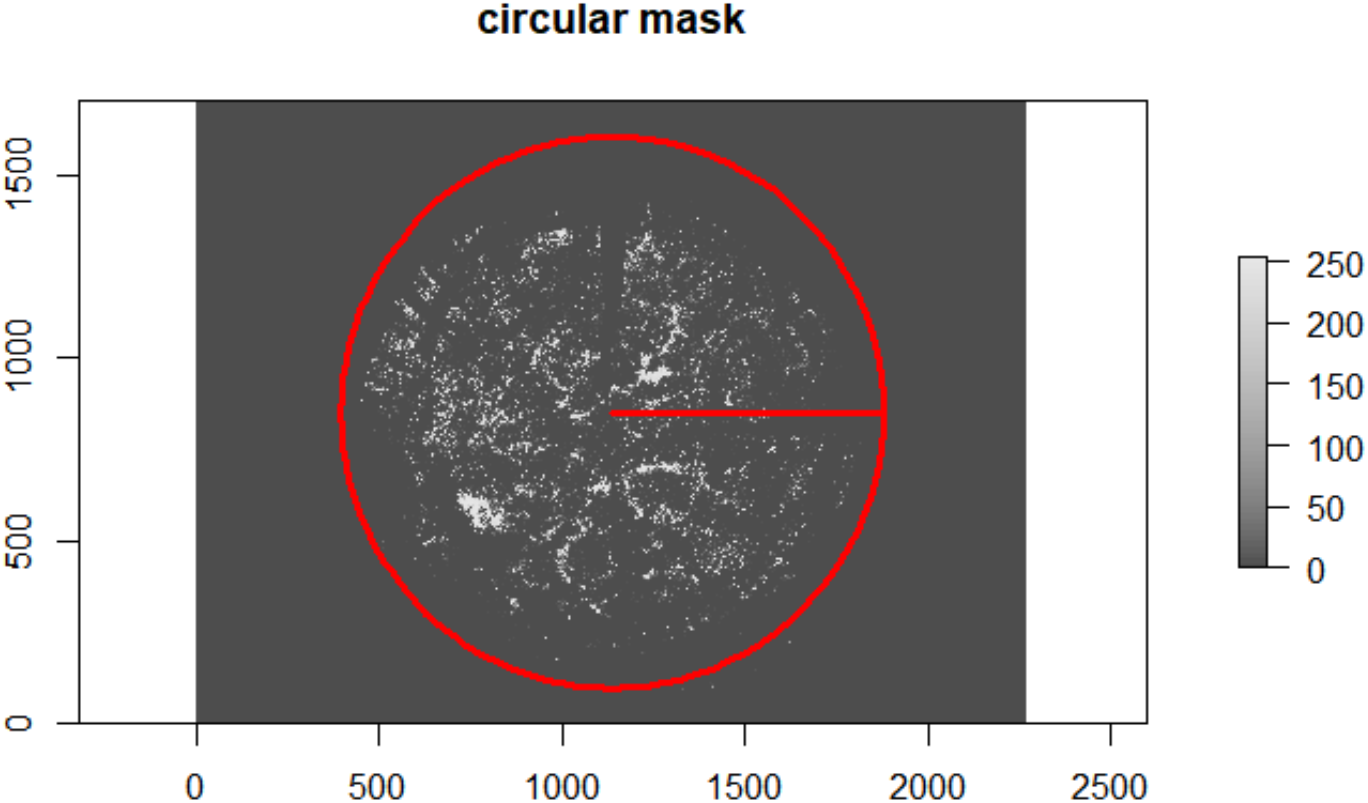
Example of a circular fisheye image uploaded in the R environment using the *import_fisheye*() function. Axis dimensions are in pixels. Red circle shows the circular mask applied, as defined by the image center (xc, yc) and radius (rc); the red line indicates the image radius from the center.

### 3.2 Create a binary image

The *binarize_fisheye()* function allows to classify the image channel imported as described above, and returns a binary raster image of canopy and gap pixels. For automated classification, the function uses the *auto_thresh()* functionality of *autothresholdr* (Schneider et al. 2012; Landini et al. 2017) to apply a single thresholding method out of 17 available ones. The default method was set to *‘Otsu’*, as it has been already proven effective in canopy images by previous studies (Pueschel et al., 2012; Grotti et al., 2020). In case of downward-looking images, which have been imported using a greenness index (e.g. GLA, GEI), we suggest using the method ‘*Percentile’*. The function also allows to set a manual threshold, using the argument ‘*manual’*; in such a case, the manual thresholding overrides the automatic classification. In case of images collected under heterogeneous sky conditions (e.g. partly cloudy, direct sunlight), the function has a ‘*zonal’* argument, which allows to divide the image in four sectors and apply the automated thresholding separately for each sector. Finally, the function has two additional arguments ‘*display’* and ‘*export’*, which respectively allows to plot the binarized classified image (Figure *4*) and export it as tiff image.

**Figure 4.**
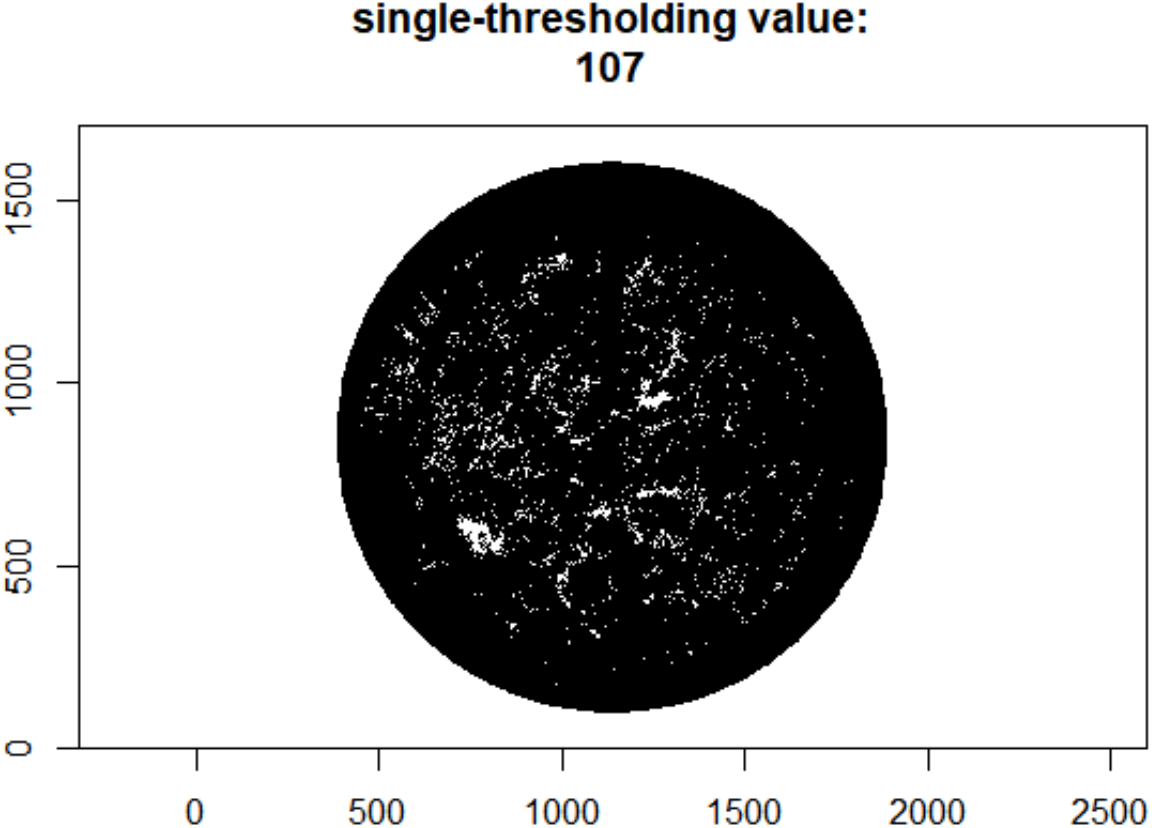
Example of a circular image, which has been classified using an automated thresholding method (‘Otsu’) using the *binarize_fisheye()* function.

### 3.3 Get angular gap fraction from binary images

The *gapfrac_fisheye()* function takes an input binary image of canopy (0) and gaps (1) and returns a dataframe of gap fraction values, grouped for an user-defined set of zenith rings (argument ‘*nrings’*) and azimuth segments (*‘nseg’*). The function uses the ‘*lens*’ argument to correct for fisheye lens distortion (namely, set the position of zenith rings based on the adjusted angular distance); it applies lens-specific projection functions, which are available for a known set of 40 fisheye lenses, which were compiled from various sources (Bourke, 2016; Pekin and Macfarlane, 2009; Schleppi et al., 2007; Thimonier et al., 2010). The general ‘equidistant’, ‘orthographic’, ‘stereographic ‘and ‘solid’ projections are also available from the *‘lens’* argument (Bourke, 2016). The complete list of available lenses for lens correction is reported in Table *2*.

**Table 2.** List of lenses available for correcting fisheye lens distorsion. The list can be screened in R environment by typing ‘list.lenses’.

| Name | Lens manufacturer | Lens model | Lens FOV |
| --- | --- | --- | --- |
| Aico-ACHIR01028B10M | Aico | ACHIR01028B10M | 245 |
| Aico-ACHIR01420B9M | Aico | ACHIR01420B9M | 183 |
| ArduCam-M25156H18 | ArduCam | M25156H18 | 180 |
| Bosch-Flexidome-7000 | Bosch | Flexidome-7000 | 180 |
| Canon-RF-5.2 | Canon | RF-5.2 | 190 |
| CanonEF-8-15 | Canon | EF-8-15 | 180 |
| DZO-VRCA | DZ Optics | VRCA | 220 |
| Entanya-HAL-200 | Entaniya | HAL-200 | 200 |
| Entanya-HAL-250 | Entaniya | HAL-250 | 250 |
| Entanya-M12-220 | Entaniya | M12-220 | 220 |
| Entanya-M12-250 | Entaniya | M12-250 | 250 |
| Entanya-M12-280 | Entaniya | M12-280 | 280 |
| Evetar-E3267A | Evetar | E3267A | 190 |
| Evetar-E3279 | Evetar | E3279 | 190 |
| Evetar-E3307 | Evetar | E3307 | 195 |
| FC-E8 | Nikon | FC-E8 | 183 |
| FC-E9 | Nikon | FC-E9 | 190 |
| iZugar-MKX13 | iZugar | MKX13 | 185 |
| iZugar-MKX19 | iZugar | MKX19 | 194 |
| iZugar-MKX200 | iZugar | MKX200 | 200 |
| iZugar-MKX22 | iZugar | MKX22 | 220 |
| Kodak-SP360 | Kodak | SP360 | 360 |
| Laowa-4 | Laowa | 4mm f/2.8 | 210 |
| Lensagon-BF10M14522S118 | Lensagon | BF10M14522S118 | 190 |
| Lensagon-BF16M220D | Lensagon | BF16M220D | 220 |
| Meike-3.5 | Meike | 3.5 mm | 220 |
| Meike-6.5 | Meike | 6.5 mm | 190 |
| Nikkor-10.5 | Nikon | 10.5 mm f/2.8 | 180 |
| Nikkor-8 | Nikon | 8 mm f/2.8 | 180 |
| Nikkor-OP10 | Nikon | 10 mm OP f/5.6 | 180 |
| Omnitech-ORIFL190-3 | Omnitech | ORIFL190-3 | 190 |
| Raynox-CF185 | Raynox | CF185 | 185 |
| Sigma-4.5 | Sigma | 4.5 mm f/2.8 | 180 |
| Sigma-8 | Sigma | 8 mm f/3.5 | 180 |
| SMTEC-SL-190 | SMTEC | SL-190 | 180 |
| Soligor | Soligor | Fisheye adapter 0.15x | 180 |
| Sunex-DSL239 | Sunex | DSL239 | 186 |
| Sunex-DSL315 | Sunex | DSL315 | 190 |
| Sunex-DSL415 | Sunex | DSL415 | 195 |
| Sunex-DSLR01 | Sunex | DSLR01 | 185 |

In addition to the number of zenith and azimuth bins, users are required to define the minimum and the maximum zenith angle for the analysis. This option increases the flexibility of the tool and allows to implement two specific camera settings, which are respectively comparable to i) LAI-2000/2200 (Licor Instruments) method and ii) the hinge-angle method. In the former, the suggested setting is to use 5 zenith rings, each 15° in size, for a zenith angle range 0°-75° (Figure *5*). In this way, the effective leaf area index (*Le*) retrieved from the subsequent *canopy_fisheye()* function is comparable to the apparent LAI derived from the LAI-2000/2200 Plant Canopy Analyzer (Chianucci et al., 2015; Ryu et al., 2010).

The hinge-angle method is based on restricted-view images centered at 1 rad (∼57.3°), as this portion of the hemisphere is insensitive to actual leaf inclination distribution of leaves (Bonhomme and Chartier, 1972). For this method, the suggested setting is to use 1 zenith ring, and a zenith angle range 55°-60° (Figure *6*).

Finally, the function has an additional arguments ‘*display’* to allows plotting the overlaid zenith and azimuth bins applied to the classified image (see examples in Figure 5 and Figure 6).

**Figure 5.**
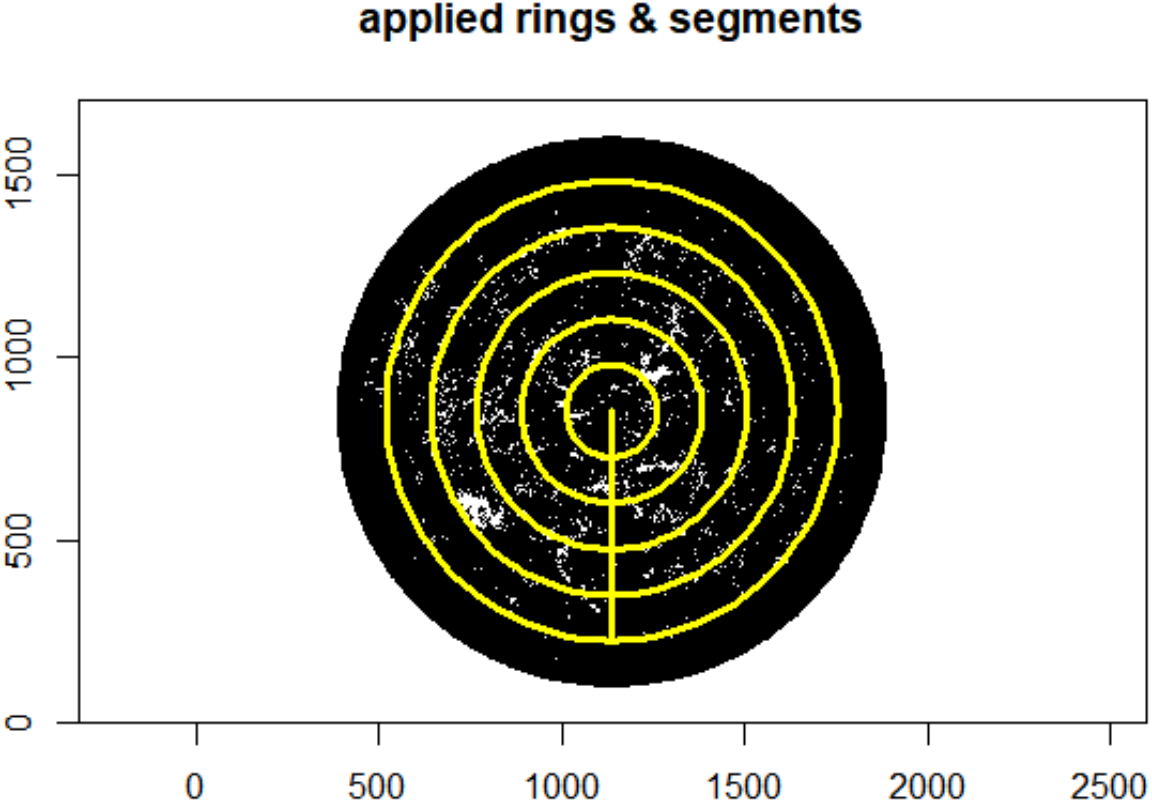
A suggested set for comparability with LAI-2000/2200 is to set 5 rings, each 15° in size. Image derived from the *gapfrac_fisheye()* function.

**Figure 6.**
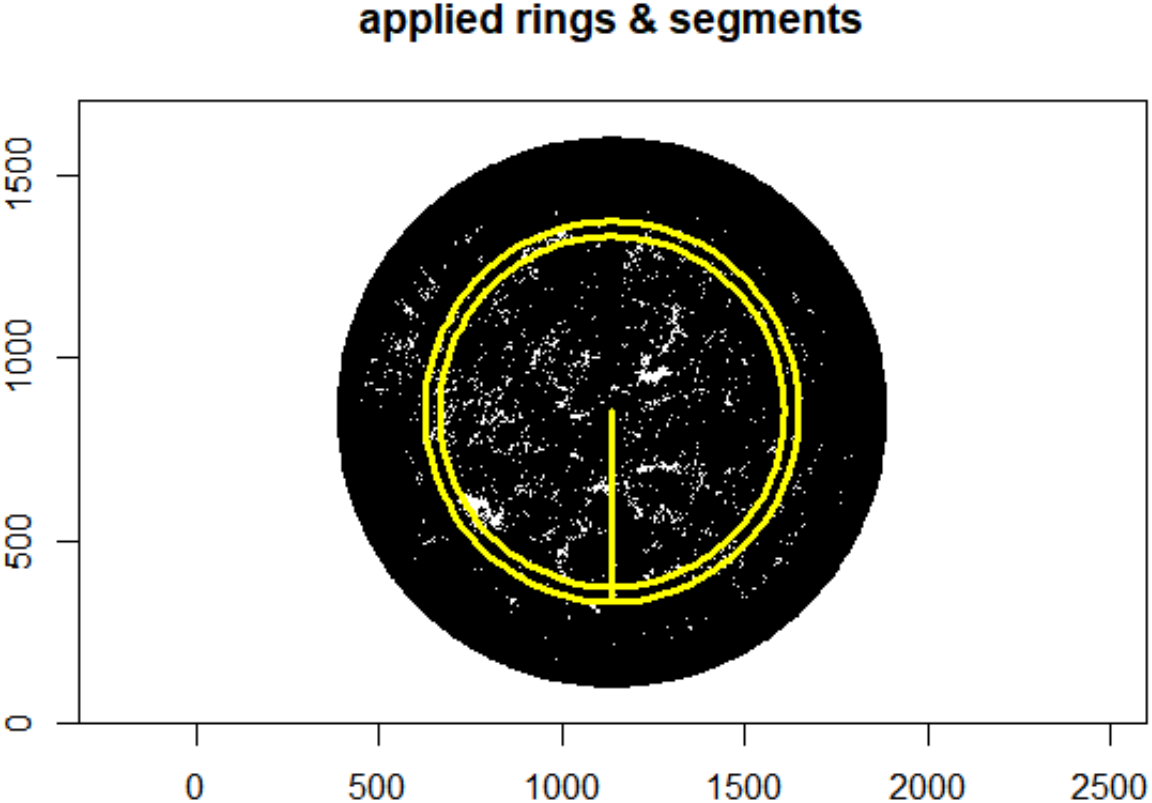
A suggested set for implementing the hinge-angle method. Image derived from the *gapfrac_fisheye()* function-

### 3.4 Retrieve canopy attributes from angular gap fraction

The function *canopy_fisheye()* applies theoretical formulas relating canopy structure to angular gap fraction. Both effective (*Le*) and actual (*L*) LAI are calculated from Miller (1967) theorem (Equation 3), by applying two different gap fraction averaging formulas (Lang and Xiang, 1986; Ryu et al., 2010):

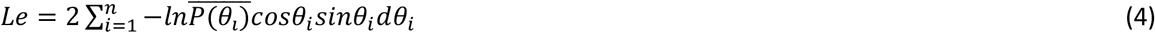

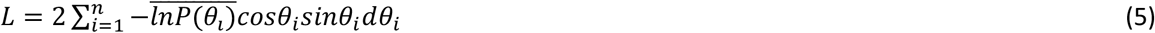

Following Ryu et al. (2010), *Le* is calculated from the logarithm of the arithmetic mean gap fraction (Equation 4), while *L* is calculated from the arithmetic mean of the logarithms of gap fraction (Equation 5), calculated for a number of zenith rings and azimuth segments. By definition *Le* ignores clumping, while *L* considers clumping at scales larger than the azimuth segment (for details, see Chianucci et al., 2019, 2015). A clumping index (LX) is then calculated from the ratio of *Le* to *L*.

Two alternative clumping indices (LXG1 and LXG2) have been calculated from ordered weighted averaging gap fraction as:

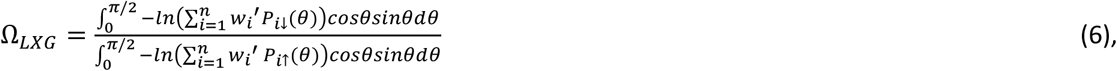

where *w*_*i*_′ is the normalized weight 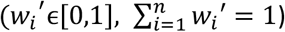 applied to decreasingly *P*_*i*↓_ and increasingly *P*_*i*↑_ ordered gap fraction. Two different monotone decreasing weighting vectors have been computed as (Ahn, 2006):

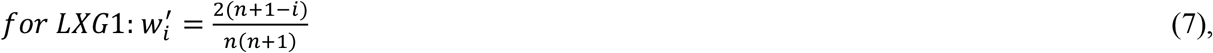

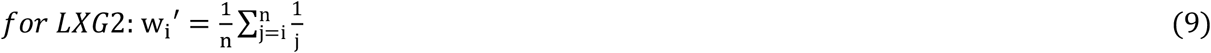

Compared with LX, LXG considers both the gap fraction distribution (as in LX), and the gap magnitude, resulting in higher clumping correction than LX. For details, see Chianucci et al. (2019).

Diffuse non-interceptance (*DIFN*), also called canopy openness, is calculated as the mean gap fraction weighted for the ring area as:

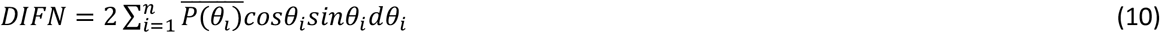

Where *cosθ*_*i*_*sinθ*_*i*_*dθ*_*i*_ is normalized to sum as 0.5, following the LAI-2000/2200 manual (*LAI-2200 Plant Canopy Analyzer Instruction Manual. LI-COR, Inc*., *Lincoln, NE*., 2012).

Additional variables calculated from the formula includes the mean leaf inclination angle (MTA.ell), which was derived by assuming an ellipsoidal foliage angle distribution:_*i*_

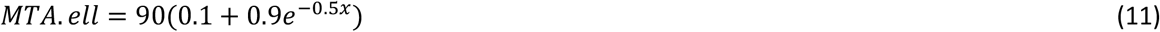

Where x is derived from a non-least square iterative procedure (Campbell, 1986). The x parameter is also provided as output of the function. It could be used to calculate the ellipsoidal extinction coefficient, as:

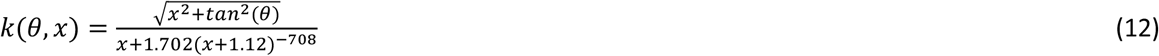

## 4. Case study

To evaluate the reliability of canopy attributes from *hemispheR*, a first test was conducted to compare canopy openness, effective (*Le*) and actual LAI (*L*), derived from the package, against values derived from WinSCANOPY (Regent Instrument, Canada) software. WinSCANOPY is a commercial software, which is often considered a reference in fisheye canopy image analysis (Chianucci and Cutini, 2013; Macfarlane et al., 2007b). The trial was performed using fisheye images collected using the Nikon Coolpix4500 equipped with the FC-E8 fisheye lens converter. Nine to 15 images have been collected in 10-12 stands, which were sampled for three years (2011-2013) in a previous study (Alivernini et al., 2018), for a total of 299 images. The stands consisted of pure deciduous forests, which are dominated by beech (*Fagus sylvatica* L.), Turkey oak (*Quercus cerris* L.) or chestnut (*Castanea sativa* Mill.), with different tree and canopy density, resulting from different silvicultural management applied. The reference LAI (as measured from littertraps) in these stands ranged from 2.6 to 7.9. Images were collected as maximum resolution JPEG and acquired close to sunrise (or sunset) under uniform sky conditions, with the aperture fixed to minimum (F5.3) and with the camera in aperture-priority (A) mode; the exposure was metered in an adjacent clearing. Subsequently, the mode was changed to manual (M) and the shutter speed was lowered by two stops in comparison to the exposure metered in the clearing (Zhang et al., 2005). A gamma correction was also applied to images prior to image processing.

Images were analyzed by setting the appropriate circular mask and lens correction in both tools, and dividing the hemisphere in 7 zenith rings, each 10° in size, and 8 azimuth segments, as proposed in another study (Chianucci and Cutini, 2013). In *hemispheR*, the ‘Otsu’ method was used to threshold images, while the method used in WinSCANOPY is not subject to scrutiny. Comparison indicated that the package reliably estimates the considered canopy attributes (Figure 7), as compared with WinSCANOPY:

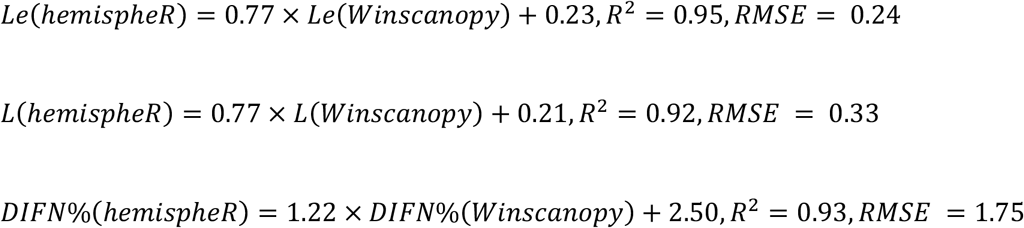

**Figure 7.**
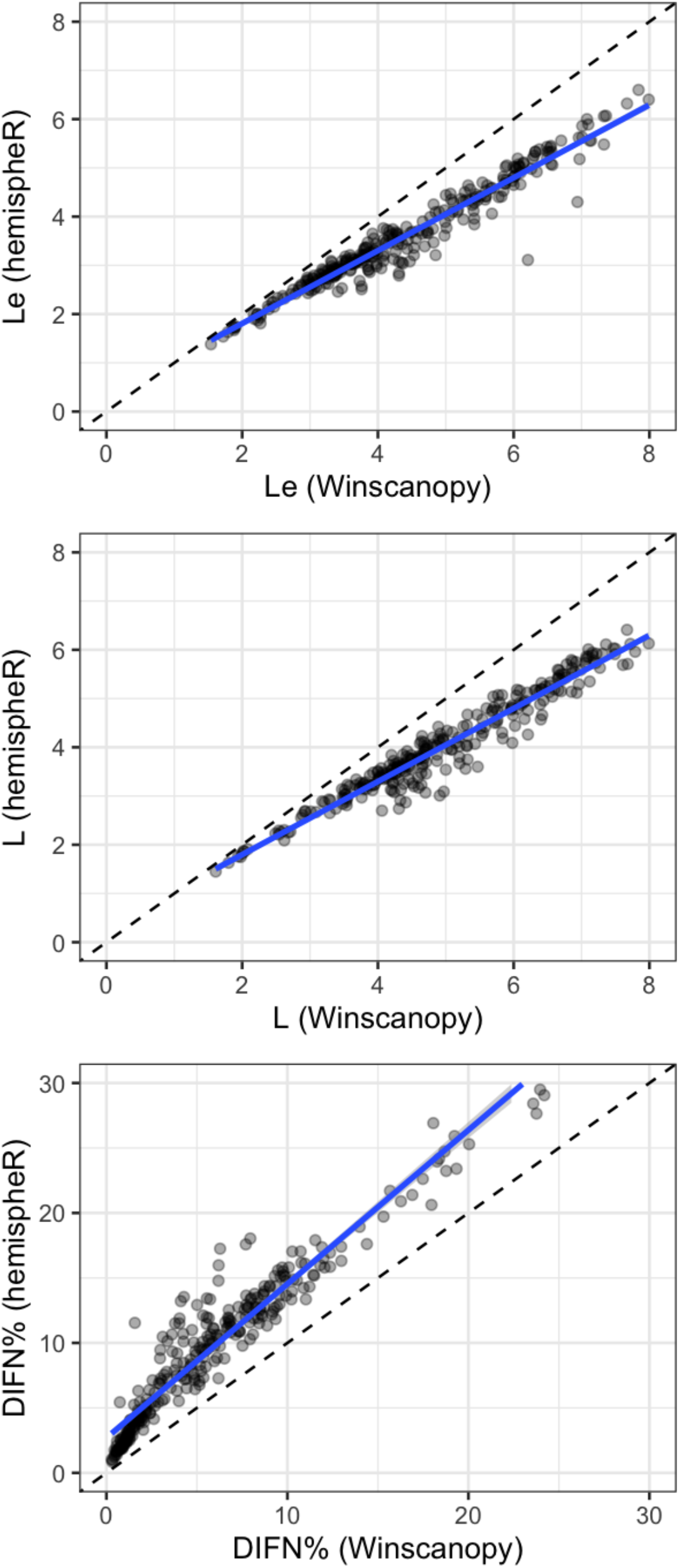
Comparison between canopy attributes derived from *hemispheR* package (y-axis) against those derived from WinSCANOPY (x-axis).

**Figure 8.**
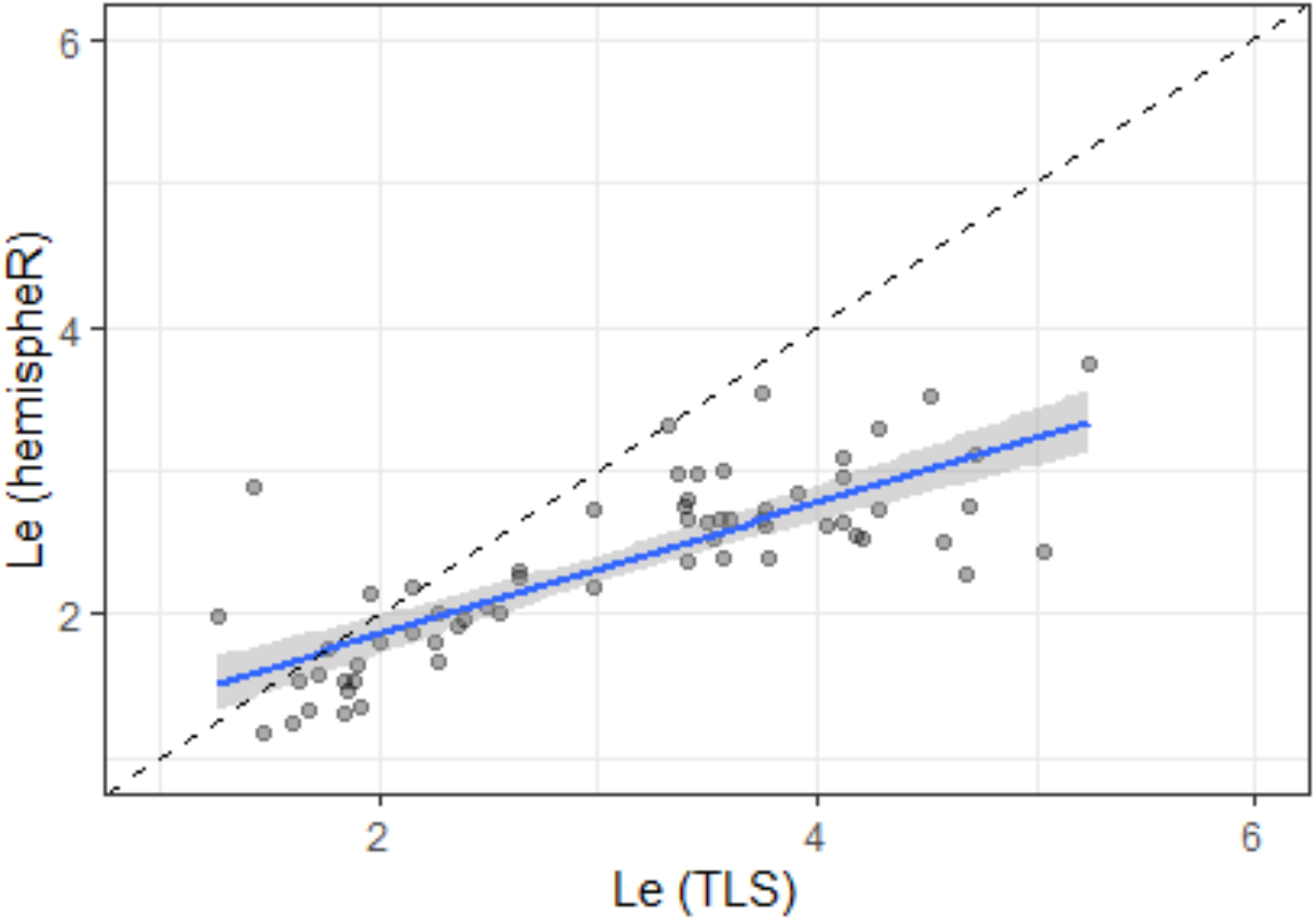
Comparison between effective leaf area index (Le) derived from *hemispheR* package (y-axis) against those derived from terrestrial laser scanning (TLS; x-axis).

Another comparison was made between fisheye images collected in seven pure beech forest stands, each 0.25 ha in size, and reference measurements obtained with a phase-shift terrestrial laser scanning; data available from a previous study (Grotti et al., 2020). In each stand, Nine DHP images and TLS scans were acquired along a grid of sampling points. TLS data were collected using a phase-shift FARO Focus 3D X130 laser scanner (Faro Technologies Inc., USA). Gap fraction was derived from TLS using a multi-thresholding method (Otsu, 1979), applied to intensity images derived from raw scan data, by setting a zenith angle range of 0-75° and 5 zenith bins, each 15° in size. *Le* was then derived using Equation 4. DHP images were collected in diffuse sky conditions with the Nikon D90 digital single lens reflex camera equipped with a Sigma 4.5 fisheye lens. Images were acquired as maximum resolution JPG and then processed in *hemispheR* using the ‘Otsu’ thresholding method, and *Le* was derived using the same number and size of zenith rings as for TLS, and the same inversion (Equation 4). Comparison indicated that *Le* from DHP significantly agreed with benchmarking values, despite a tendency to underestimate *Le* compared with TLS.

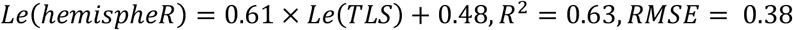

The underestimation of *Le* in DHP was in line with the observed ability of the active sensor to detect smaller canopy elements, particularly at increasing zenith angle (longer optical distance), due to the vastly higher resolution of laser scanning data (Grotti et al., 2020).

## Discussion and conclusions

A recent review by Atkins et al., (2022) states that “the R programming language offers a viable and rich option for forestry and forest ecology research with a diverse and plentiful user community and development base. Further adoption of R in forest-research could increase the visibility and reproducibility of research, while also increasing productivity”. From this perspective, we believe that *hemispheR* offers a potentially-relevant tool to further make fisheye photography a simple and accessible method, making the process robust, automated, and transparent. In our view, the key points of the package are:

- the embedding of *raster* functionality, which allows importing and processing each kind of raster (i.e., pixel matrix) image, with great integration with other R image processing packages;
- the sequentiality of functions, which makes the package simple to use, while ensuring code conciseness and clarity, and suitable for pipeline integration;
- the possibility to process both circular and fullframe fisheye images, which can significantly extend the camera and lens types available for hemispherical image analysis, including smartphones camera and lenses (Andis, 2022; Chianucci, 2019);

With reference on package limitations, *hemispheR* has not implemented specific features for developing raw imagery, which may represent a valuable solution for reducing the exposure-dependency of fisheye images (Hwang et al., 2016; Macfarlane et al., 2014). However, such option is already available in some R packages such as *‘adimpro’* (Polzehl and Tabelow, 2007), which can be used in combination with the *hemispheR* package. With reference on thresholding methods, only single thresholding values are considered in the package; future implementations can consider also dual thresholding (Macfarlane, 2011), or object-oriented classifications; even in this case, the package can be used in combination with other packages already performing such tasks such as ‘*caiman’* (Díaz et al., 2021). In addition, as the package is mostly focused on forest canopies (i.e., upward facing camera orientation), the development of downward looking solutions (e.g. use of greenness indices) are still at an experimental stage, and need to be validated against benchmark measurements. In this line, we believe that hosting the package in a Git repository will further support development of *hemispheR*, through either collaborative coding or forking projects.

## Package availability

The package hemispherR can be installed in R using devtools (Wickham et al., 2021) by typing: devtools::install_git(“https://gitlab.com/fchianucci/hemispheR”).

## References

Ahn, B.S., 2006. On the properties of OWA operator weights functions with constant level of orness. IEEE Transactions on Fuzzy Systems 14, 511–515. https://doi.org/10.1109/TFUZZ.2006.876741

Alivernini, A., Fares, S., Ferrara, C., Chianucci, F., 2018. An objective image analysis method for estimation of canopy attributes from digital cover photography. Trees 32, 713–723. https://doi.org/10.1007/s00468-018-1666-3

Anderson, M., 1964. Studies of the Woodland Light Climate: I. The Photographic Computation of Light Conditions. Journal of Ecology 52, 27–41. https://doi.org/10.2307/2257780

Andis, A.A., 2022. Estimation of forest canopy structure and understory light using spherical panorama images from smartphone photography. Forestry 95, 38–48.

Atkins, J.W., Stovall, A.E.L., Alberto Silva, C., 2022. Open-Source tools in R for forestry and forest ecology. Forest Ecology and Management 503, 119813. https://doi.org/10.1016/j.foreco.2021.119813

Bachelot, B., 2016. Sky: Canopy Openness Analyzer Package.

Bonhomme, R., Chartier, P., 1972. The interpretation and automatic measurement of hemispherical photographs to obtain sunlit foliage area and gap frequency. Israel J. Agric. Res 22, 53–61.

Bourke, P., 2016. Fisheye lens correction.

Breda, N.J.J., 2003. Ground-based measurements of leaf area index: a review of methods, instruments and current controversies. Journal of Experimental Botany 54, 2403–2417. https://doi.org/10.1093/jxb/erg263

Brown, L.A., Ogutu, B.O., Dash, J., 2020. Tracking forest biophysical properties with automated digital repeat photography: A fisheye perspective using digital hemispherical photography from below the canopy. Agricultural and Forest Meteorology 287, 107944. https://doi.org/10.1016/j.agrformet.2020.107944

Campbell, G.S., 1986. Extinction coefficients for radiation in plant canopies calculated using an ellipsoidal inclination angle distribution. Agricultural and Forest Meteorology 36, 317–321. https://doi.org/10.1016/0168-1923(86)90010-9

Chianucci, F., 2019. An overview of in situ digital canopy photography in forestry. Can. J. For. Res. 227–242. https://doi.org/10.1139/cjfr-2019-0055

Chianucci, F., Cutini, A., 2013. Estimation of canopy properties in deciduous forests with digital hemispherical and cover photography. Agricultural and Forest Meteorology 168, 130–139. https://doi.org/10.1016/j.agrformet.2012.09.002

Chianucci, F., Cutini, A., 2012. Digital hemispherical photography for estimating forest canopy properties: current controversies and opportunities. iForest 5, 290–295. https://doi.org/10.3832/ifor0775-005

Chianucci, F., Macfarlane, C., Pisek, J., Cutini, A., Casa, R., 2015. Estimation of foliage clumping from the LAI-2000 Plant Canopy Analyzer: effect of view caps. Trees 29, 355–366. https://doi.org/10.1007/s00468-014-1115-x

Chianucci, F., Salvati, L., Giannini, T., Chiavetta, U., Corona, P., Cutini, A., 2016. Long-term response to thinning in a beech (Fagus sylvatica L.) coppice stand under conversion to high forest in Central Italy. Silva Fenn. 50. https://doi.org/10.14214/sf.1549

Chianucci, F., Zou, J., Leng, P., Zhuang, Y., Ferrara, C., 2019. A new method to estimate clumping index integrating gap fraction averaging with the analysis of gap size distribution. Can. J. For. Res. 49, 471– 479. https://doi.org/10.1139/cjfr-2018-0213

Decan, A., Mens, T., Claes, M., Grosjean, P., 2015. On the Development and Distribution of R Packages: An Empirical Analysis of the R Ecosystem. ECSAW ‘15: Proceedings of the 2015 European Conference on Software Architecture Workshops.

Díaz, G.M., Negri, P.A., Lencinas, J.D., 2021. Toward making canopy hemispherical photography independent of illumination conditions: A deep-learning-based approach. Agricultural and Forest Meteorology 296, 108234. https://doi.org/10.1016/j.agrformet.2020.108234

Evans, G.C., Coombe, D.E., 1959. Hemisperical and Woodland Canopy Photography and the Light Climate. Journal of Ecology 47, 103–113. https://doi.org/10.2307/2257250.

Fang, H., Baret, F., Plummer, S., Schaepman-Strub, G., 2019. An Overview of Global Leaf Area Index (LAI): Methods, Products, Validation, and Applications. Rev. Geophys. 57, 739–799. https://doi.org/10.1029/2018RG000608

Frazer, G.W., Canham, C.D., Lertzman, 1999. Gap Light Analyzer (GLA), Version 2.0: Imaging software to extract canopy structure and gap light transmission indices from true-colour fisheye photographs, users manual and program documentation.

Glatthorn, J., Beckschäfer, P., 2014. Standardizing the Protocol for Hemispherical Photographs: Accuracy Assessment of Binarization Algorithms. PLoS ONE 9, e111924. https://doi.org/10.1371/journal.pone.0111924

Gonsamo, A., Walter, J.-M.N., Pellikka, P., 2011. CIMES: A package of programs for determining canopy geometry and solar radiation regimes through hemispherical photographs. Computers and Electronics in Agriculture 79, 207–215. https://doi.org/10.1016/j.compag.2011.10.001

Grotti, M., Calders, K., Origo, N., Puletti, N., Alivernini, A., Ferrara, C., Chianucci, F., 2020. An intensity, image-based method to estimate gap fraction, canopy openness and effective leaf area index from phase-shift terrestrial laser scanning. Agricultural and Forest Meteorology 280, 107766. https://doi.org/10.1016/j.agrformet.2019.107766

Hijmans, R.J., 2021. raster: Geographic Data Analysis and Modeling. R package version 3.4-10.

Hwang, Y., Ryu, Y., Kimm, H., Jiang, C., Lang, M., Macfarlane, C., Sonnentag, O., 2016. Correction for light scattering combined with sub-pixel classification improves estimation of gap fraction from digital cover photography. Agricultural and Forest Meteorology 222, 32–44. https://doi.org/10.1016/j.agrformet.2016.03.008

Jonckheere, I., Fleck, S., Nackaerts, K., Muys, B., Coppin, P., Weiss, M., Baret, F., 2004. Review of methods for in situ leaf area index determination. Agricultural and Forest Meteorology 121, 19–35. https://doi.org/10.1016/j.agrformet.2003.08.027

Kašpar, V., Hederová, L., Macek, M., Müllerová, J., Prošek, J., Surový, P., Wild, J., Kopecký, M., 2021. Temperature buffering in temperate forests: Comparing microclimate models based on ground measurements with active and passive remote sensing. Remote Sensing of Environment 263, 112522. https://doi.org/10.1016/j.rse.2021.112522

LAI-2200 Plant Canopy Analyzer Instruction Manual, 2012.. LI-COR, Inc., Lincoln, NE.

Landini, G., Randell, D.A., Fouad, S., Galton, A., 2017. Automatic thresholding from the gradients of region boundaries: AUTOMATIC THRESHOLDING. Journal of Microscopy 265, 185–195. https://doi.org/10.1111/jmi.12474

Lang, A.R.G., Xiang, Y., 1986. Estimation of leaf area index from transmission of direct sunlight in discontinuous canopies. Agricultural and forest Meteorology 37, 229–243. https://doi.org/10.1016/0168-1923(86)90033-X

Lang, M., Nilson, T., Kuusk, A., Pisek, J., Korhonen, L., Uri, V., 2017. Digital photography for tracking the phenology of an evergreen conifer stand. Agricultural and Forest Meteorology 246, 15–21. https://doi.org/10.1016/j.agrformet.2017.05.021

Liu, J., Pattey, E., 2010. Retrieval of leaf area index from top-of-canopy digital photography over agricultural crops. Agricultural and Forest Meteorology 150, 1485–1490. https://doi.org/10.1016/j.agrformet.2010.08.002

Louhaichi, M., Borman, M.M., Johnson, D.E., 2008. Spatially Located Platform and Aerial Photography for Documentation of Grazing Impacts on Wheat. Geocarto International 16, 65–70. https://doi.org/10.1080/10106040108542184

Macfarlane, C., 2011. Classification method of mixed pixels does not affect canopy metrics from digital images of forest overstorey. Agricultural and Forest Meteorology 151, 833–840. https://doi.org/10.1016/j.agrformet.2011.01.019

Macfarlane, C., Coote, M., White, D.A., Adams, M.A., 2000. Photographic exposure affects indirect estimation of leaf area in plantations of Eucalyptus globulus Labill. Agricultural and Forest Meteorology 100, 155– 168. https://doi.org/10.1016/S0168-1923(99)00139-2

Macfarlane, C., Grigg, A., Evangelista, C., 2007a. Estimating forest leaf area using cover and fullframe fisheye photography: Thinking inside the circle. Agricultural and Forest Meteorology 146, 1–12. https://doi.org/10.1016/j.agrformet.2007.05.001

Macfarlane, C., Hoffman, M., Eamus, D., Kerp, N., Higginson, S., McMurtrie, R., Adams, M., 2007b. Estimation of leaf area index in eucalypt forest using digital photography. Agricultural and Forest Meteorology 143, 176–188. https://doi.org/10.1016/j.agrformet.2006.10.013

Macfarlane, C., Ryu, Y., Ogden, G.N., Sonnentag, O., 2014. Digital canopy photography: Exposed and in the raw. Agricultural and Forest Meteorology 197, 244–253. https://doi.org/10.1016/j.agrformet.2014.05.014

Miller, J., 1967. A formula for average foliage density. Australian Journal of Botany 15, 141–144.

Otsu, N., 1979. Threshold selection method from gray-level histograms. EEE transactions on systems, man, and cybernetics 62–66.

Pekin, B., Macfarlane, C., 2009. Measurement of Crown Cover and Leaf Area Index Using Digital Cover Photography and Its Application to Remote Sensing. Remote Sensing 1, 1298–1320. https://doi.org/10.3390/rs1041298

Polzehl, J., Tabelow, K., 2007. Adaptive Smoothing of Digital Images: The R Package adimpro. Journal of Statistical Software 19, 17. https://doi.org/10.18637/jss.v019.i01

Pueschel, P., Buddenbaum, H., Hill, J., 2012. An efficient approach to standardizing the processing of hemispherical images for the estimation of forest structural attributes. Agricultural and Forest Meteorology 160, 1–13. https://doi.org/10.1016/j.agrformet.2012.02.007

Ryu, Y., Nilson, T., Kobayashi, H., Sonnentag, O., Law, B.E., Baldocchi, D.D., 2010. On the correct estimation of effective leaf area index: Does it reveal information on clumping effects? Agricultural and Forest Meteorology 150, 463–472. https://doi.org/10.1016/j.agrformet.2010.01.009

Schleppi, P., Conedera, M., Sedivy, I., Thimonier, A., 2007. Correcting non-linearity and slope effects in the estimation of the leaf area index of forests from hemispherical photographs. Agricultural and Forest Meteorology 144, 236–242. https://doi.org/10.1016/j.agrformet.2007.02.004

Sercu, B.K., Baeten, L., van Coillie, F., Martel, A., Lens, L., Verheyen, K., Bonte, D., 2017. How tree species identity and diversity affect light transmittance to the understory in mature temperate forests. Ecology and Evolution 24, 10861–10870.

Skelly, D.K., Halverson, M.A., Freidenburg, L.K., Urban, M.C., 2005. Canopy closure and amphibian diversity in forested wetlands. Wetlands Ecol Manage 13, 261–268. https://doi.org/10.1007/s11273-004-7520-y

er Steege, H., 2018. Hemiphot. R: Free R scripts to analyse hemispherical photographs for canopy openness, leaf area index and photosynthetic active radiation under forest canopies.

Thimonier, A., Sedivy, I., Schleppi, P., 2010. Estimating leaf area index in different types of mature forest stands in Switzerland: a comparison of methods. Eur J Forest Res 129, 543–562. https://doi.org/10.1007/s10342-009-0353-8

Thomas, S.C., Halpern, C.B., Falk, D.A., Liguori, D.A., Austin, K.A., 1999. PLANT DIVERSITY IN MANAGED FORESTS: UNDERSTORY RESPONSES TO THINNING AND FERTILIZATION. Ecological Applications 9, 864– 879. https://doi.org/10.1890/1051-0761(1999)009[0864:PDIMFU]2.0.CO;2

Weiss, Marie, Baret, F., 2017. CAN_EYE V6. 4.91 user manual.

Wickham, H., Hester, J., Chang, W., Bryan, J., 2021. devtools: Tools to Make Developing R Packages Easier}.

Woebbecke, D.M., Meyer, G.E., Von Bargen, K., Mortensen, D.A., 1995. Color Indices for Weed Identification Under Various Soil, Residue, and Lighting Conditions. Transactions of the ASAE. https://doi.org/10.13031/2013.27838

Yan, G., Hu, R., Luo, J., Weiss, M., Jiang, H., Mu, X., Xie, D., Zhang, W., 2019. Review of indirect optical measurements of leaf area index: Recent advances, challenges, and perspectives. Agricultural and Forest Meteorology 265, 390–411. https://doi.org/10.1016/j.agrformet.2018.11.033

Zhang, Y., Chen, J.M., Miller, J.R., 2005. Determining digital hemispherical photograph exposure for leaf area index estimation. Agricultural and Forest Meteorology 133, 166–181. https://doi.org/10.1016/j.agrformet.2005.09.009

Zhao, K., Ryu, Y., Hu, T., Garcia, M., Li, Y., Liu, Z., Londo, A., Wang, C., 2019. How to better estimate leaf area index and leaf angle distribution from digital hemispherical photography? Switching to a binary nonlinear regression paradigm. Methods in Ecology and Evolution 10, 1864–1874. https://doi.org/10.1111/2041-210X.13273

